# Examining craniofacial variation among crispant and mutant zebrafish models of human skeletal diseases

**DOI:** 10.1101/2022.08.18.504429

**Authors:** Kelly M. Diamond, Abigail E. Burtner, Daanya Siddiqui, Kurtis Alvarado, Sanford L. Leake, Sara Rolfe, Chi Zhang, Ronald Y. Kwon, A. Murat Maga

## Abstract

Genetic diseases affecting the skeletal system present with a wide range of symptoms that make diagnosis and treatment difficult. Genome-wide association and sequencing studies have identified genes linked to human skeletal diseases. Gene editing of zebrafish models allows researchers to further examine the link between genotype and phenotype, with the long-term goal of improving diagnosis and treatment. While current automated tools enable rapid and in-depth phenotyping of the axial skeleton, characterizing the effects of mutations on the craniofacial skeleton has been more challenging. The objective of this study was to evaluate a semi-automated screening tool can be used to quantify craniofacial variations in zebrafish models using four genes that have been associated with human skeletal diseases (*meox1, plod2, sost*, and *wnt16*) as test cases. We used traditional landmarks to ground truth our dataset and pseudolandmarks to quantify variation across the 3D cranial skeleton between the groups (somatic crispant, germline mutant, and control fish). The proposed pipeline identified variation between the crispant or mutant fish and control fish for four genes. Variation in phenotypes parallel human craniofacial symptoms for two of the four genes tested. This study demonstrates the potential as well as the limitations of our pipeline as a screening tool to examine multi-dimensional phenotypes associated with the zebrafish craniofacial skeleton.

## Introduction

The Nosology Committee of the International Skeletal Dysplasia Society currently recognizes 461 skeletal disorders and 437 genes that are associated with 425 of these disorders (Mortier et al., 2019). As these associations are identified, methods such as Next Generation Sequencing and genetic modification of model organisms are used to parse out the underlying genetic mechanisms of skeletal diseases. CRISPR-modified mouse and zebrafish models have been used to understand the mechanisms for skeletal diseases such as sclerosteosis, osteoporosis, and osteogenesis imperfecta, as well as more rare variations of these disorders (Garg et al., 2022; Kaya et al., 2022; Kwon et al., 2019). Using model organisms to understand the genetic mechanism of these disorders presents opportunities to develop and test potential treatments (Kague and Karasik, 2022). However, for these models to be useful, the homologous genes in model organisms must present observable phenotypes that parallel the symptoms of human diseases. Current best practices use measures such as tissue mineral density and bone thickness to identify phenotypes that are common between model organisms and human diseases. In this study we present a semiautomatic pipeline to quantitatively assess the craniofacial phenotype of adult zebrafish.

The zebrafish genome is identical to 71% of the human genome, with 47% of human genes having a one-to-one correspondence with zebrafish genes (Howe et al., 2013). Additionally, as teleosts, zebrafish share numerous conserved features of their skeletal system and ossification mechanisms with mammals (Dietrich et al., 2021). While mice, as mammals, do share a larger percentage of their genome with humans, zebrafish present several advantages for use as a model system for studying skeletal disease. A benefit of using zebrafish is that zebrafish eggs are fertilized, and embryos develop externally. By alleviating the need for embryo transfer for CRISPR-induced gene editing as in mouse, this facilitates rapid and efficient mutagenesis in zebrafish. Second, zebrafish are transparent through early development, facilitating the visual inspection of developing skeletal structures in live embryos and larvae. Finally, the zebrafish skeletal system develops rapidly, with ossification beginning 3-4 days post fertilization and the skeletal maturity reached by 2-4 months (Dietrich et al., 2021). For these reasons, zebrafish are increasingly used in studies of skeletal disorders (Busse et al., 2020; Kwon et al., 2019).

Much of the past work on the development of semi-automated phenotyping methods in adult zebrafish has focused on the axial skeleton (Hur et al., 2017; Watson et al., 2020). In contrast to the craniofacial skeleton, the meristic and segmented nature of the axial skeleton makes automating the quantitative phenotyping more straight-forward (Hur et al., 2017). Skeletal remodeling, which is often affected by skeletal disease, is highly dependent on bone loading, hence focusing on aspects of the skeleton that have a greater proportion of skeletal muscles due to their use for locomotion is logical (Dietrich et al., 2021). However, the craniofacial skeleton is influenced by both developmental patterning and mechanically induced remodeling (Conith et al., 2019). Humans and zebrafish craniofacial skeletons differ in the number of bones; but the embryonic origins of these bones are homologous. In zebrafish, the anterior and lateral bones of the craniofacial skeleton are derived from neural crest ectoderm, while the posterior portions of the cranial skeleton are derived from mesoderm (Galea et al., 2021; Kague et al., 2012). Additionally, zebrafish have four major bone types across the cranial skeleton (acellular compact, cellular compact, tubular, and spongy), which differ in their formation (Weigele and Franz-Odendaal, 2016). Assessing the phenotype of bones that use different methods of ossification in crispant and mutant zebrafish could provide more insight into the underlying mechanisms of skeletal diseases.

In this study, we build on a previously developed semiautomated pipeline (Diamond et al., 2022) to identify areas of craniofacial variation in four different zebrafish models of skeletal disease: *plod2, meox1, sost* and *wnt16*. Zebrafish models of *plod2* are associated with Bruck Syndrome, a rare form of Osteogenesis Imperfecta (Lv et al., 2018). Mutations in *meox1* are linked to Klippel–Feil syndrome, which presents with the fusion of cervical vertebrae (Dauer et al., 2018). *sost* mutations have been linked to sclerosteosis, or hyperosteosis of the skull and long bones (Hamersma et al., 2003). Finally, *wnt16* is associated with osteoporosis (Medina-Gomez et al., 2012). We used three somatic crispant models (*plod2, meox1*, and *wnt16*) and two germline mutant models (*sost* and *wnt16*) to examine the extent to which this pipeline could be used to identify areas of variation across the zebrafish skull. For each model we examined mutant or crispant fish and control siblings to test for the effects of each knockout on craniofacial phenotype. We predicted that craniofacial phenotypes for each gene will reflect human disease phenotypes as these models have all successfully replicated human disease phenotypes associated with axial skeleton. In addition to running individual analyses, we also examined shape differences across the entire dataset and made two predictions about the combined analysis. We first predicted that control fish from all experiments should be grouped closer together in morphospace, as all control fish should have more similar genetic backgrounds than any control fish has to the mutant groups. Our second prediction was that the mutant lines, especially crispant fish, should be closer in morphospace to the control fish from their respective experiment, as controls are from the same clutch. By examining within and among these different models we aim to test the effectiveness of this pipeline for broad use in phenotyping the craniofacial skeleton of a model organism that is heavily used in studies of developmental and evolutionary biology.

## Materials & Methods

The dataset for this study includes somatic crispant and/or germline mutant zebrafish (*Danio rerio*) lines of four genes (*plod2, meox1, sost*, and *wnt16*) whose human homologs have previously been associated with human skeletal disease. We used clutches that included crispant fish and their wildtype siblings for *plod2* (10 crispant; 11 control fish), *meox1* (11 crispant; 11 control fish), and *wnt16* (22 *wnt16* crispant; 18 control fish). We also included germline mutant lines for *sost* (7 mutants; 9 heterozygotes; 8 control fish) and *wnt16* (5 mutants; 6 heterozygotes; 3 control fish). Generation of crispants for *plod2* and *wnt16* was previously described (Watson et al., 2020; Watson et al., 2021); Generation of crispants for *meox1* was performed using the methods described in Watson et al. 2020. In this case, Cas9:gRNAs targeting two distinct regions (gRNA1: GTCAGGAGTCCTCATTCGGG; gRNA2: GGTTGACTGCGATCTCGTAG) were injected. Germline mutants for *wnt16* (*wnt16*^*w1001*^) were previously described in Watson et al. 2021. Germline mutants for *sost* (Tg[sost:NTR-GFP]^w216^) were previously described (Thomas and Raible, 2019). Mutant alleles for both *wnt16* and *sost* are both predicted to be null (Thomas and Raible, 2019; Watson et al., 2021). Studies were conducted in mixed sex animals. All fish were sacrificed at approximately 90 days post fertilization and scanned using a vivaCT40 MicroCT scanner (Scanco Medical, Switzerland), with 21micron isotropic voxel resolution, 55kVp, 145mA, 1024 samples, 500proj/180°, 200ms integration time. All studies were performed on an approved protocol in accordance with the University of Washington Institutional Animal Care and Use Committee (IACUC# 4306-01).

Before beginning our individual assessments of each gene pipeline, we first examined the digitizing accuracy of the team by having all digitizers (authors AB, DS, KA, KD, SL) manually landmark 21 points on the *plod2* zebrafish skulls (see Table S1 for specific points) using the Markups module of 3D Slicer (Fedorov et al., 2012). We analyzed these landmarks using the geomorph package in R (Adams and Otárola-Castillo, 2013). We first ran a combined Generalized Procrustes Analysis (GPA) and tested if overall shape varied among digitizers using Procrustes ANOVA (Goodall, 1991). We then performed a Principal Components Analysis (PCA) on the coordinates from the GPA to examine if there were any specific issues with our digitizing error.

For each gene, we first compared the 21 manual landmarks to establish the baseline set of differences among groups. With the exception of *plod2*, in which the average of all landmarks was used as our manual gold standard, all genes were landmarked by a single author. Next, we transferred the set of 308 pseudolandmark points using the normative atlas of wildtype individuals that we previously published (Diamond et al., 2022) using the ALPACA module (Porto et al., 2021) of the SlicerMorph extension (Rolfe et al., 2021) in 3D Slicer (Fedorov et al., 2012). Pseudolandmark points were created using the PseudoLMGenerator module of the SlicerMorph package in 3D Slicer. Our sample sizes for the pseudolandmarks were smaller for the *plod2* (10 control fish), *wnt16* crispant (20 crispant, 15 control fish), and *wnt16* germline mutant (4 mutant fish) groups due to challenges with converting the microCT scans into the 3D mesh files necessary for our pseudolandmark pipeline.

For each gene and dataset (manual landmarks versus pseudolandmarks), we independently tested if overall skull shape varied between mutant and control fish using a Procrustes ANOVA model (shape ∼ group), with group defined as mutant or control, and shape defined as the Procrustes coordinates from the GPA. If shape differences were found between control and mutant fish, we ran a PCA. For all principal components (PC) that explained more than 2% of the variation in the PCA, we tested if mutant and control fish varied along those PCs using a Procrustes ANOVAs model (PC ∼ group), with group again defined as mutant or control, and PC defined as the PC scores from the PCA. Both the manual and pseudolandmark points were analyzed using the geomorph package in R (Adams and Otárola-Castillo, 2013). Because our pseudolandmark points were bilaterally symmetrical, we also ran a symmetry analysis for each gene using the geomorph package in R (Adams and Otárola-Castillo, 2013). We supply an example script for the full analysis of a single gene in Appendix 1.

When mutant and control fish varied along a PC in any of our analyses, we visualized the shape differences across the PC with heatmaps. To create the heatmaps, we first created models of average shape as well as the minimum and maximum shape for each PC using the morpho package in R (Schlager, 2017). We then used a custom python script (Appendix 2) to extract the mesh distances over only the areas where we had pseudolandmark points. Heatmaps of these distances were then created using the Models module of 3D Slicer.

In addition to individual group comparisons, we also performed combined analyses for both the 21 manual and 308 pseudolandmark points. Both datasets were analyzed with the same methods as the individual analyses stated above. For the manual landmark comparisons, we also wanted to ensure any patterns we found were not due to differences between the individuals who landmarked the different datasets. To do this we used the MALPACA module (Zhang et al., 2022) of the SlicerMorph extension (Rolfe et al., 2021) in 3DSlicer (Fedorov et al., 2012). We used one control fish from each gene as templates and then used the median of the grouped outputs for each fish in the dataset.

## Results & Discussion

### Groupwise differences

In our quality control experiment we had all individuals who digitized the various datasets in this study landmark the same dataset (*plod2* fish and control siblings). When this dataset was analyzed, we found overall shape varied modestly among digitizing individuals (F=1.440, z=1.751, p=0.046). We argue this result supports our recommendation to use a pseudolandmarking approach, as the geometrically homologous points are not susceptible to variation among different digitizers. However, we also acknowledge the need to ground truth our dataset, so while we recognize some variation in our manual landmarks are due to human-induced variation, we have chosen to include a groupwise comparison of manual landmarks to show how overall patterns are similar between the manual and pseudolandmark datasets. We also tested if any of the individual Principal Components (PC) vary by digitizer and found PC3, PC5, and PC9 differ, however this variation is minimal (Figure 1A; Table S2).

**Figure 1.**
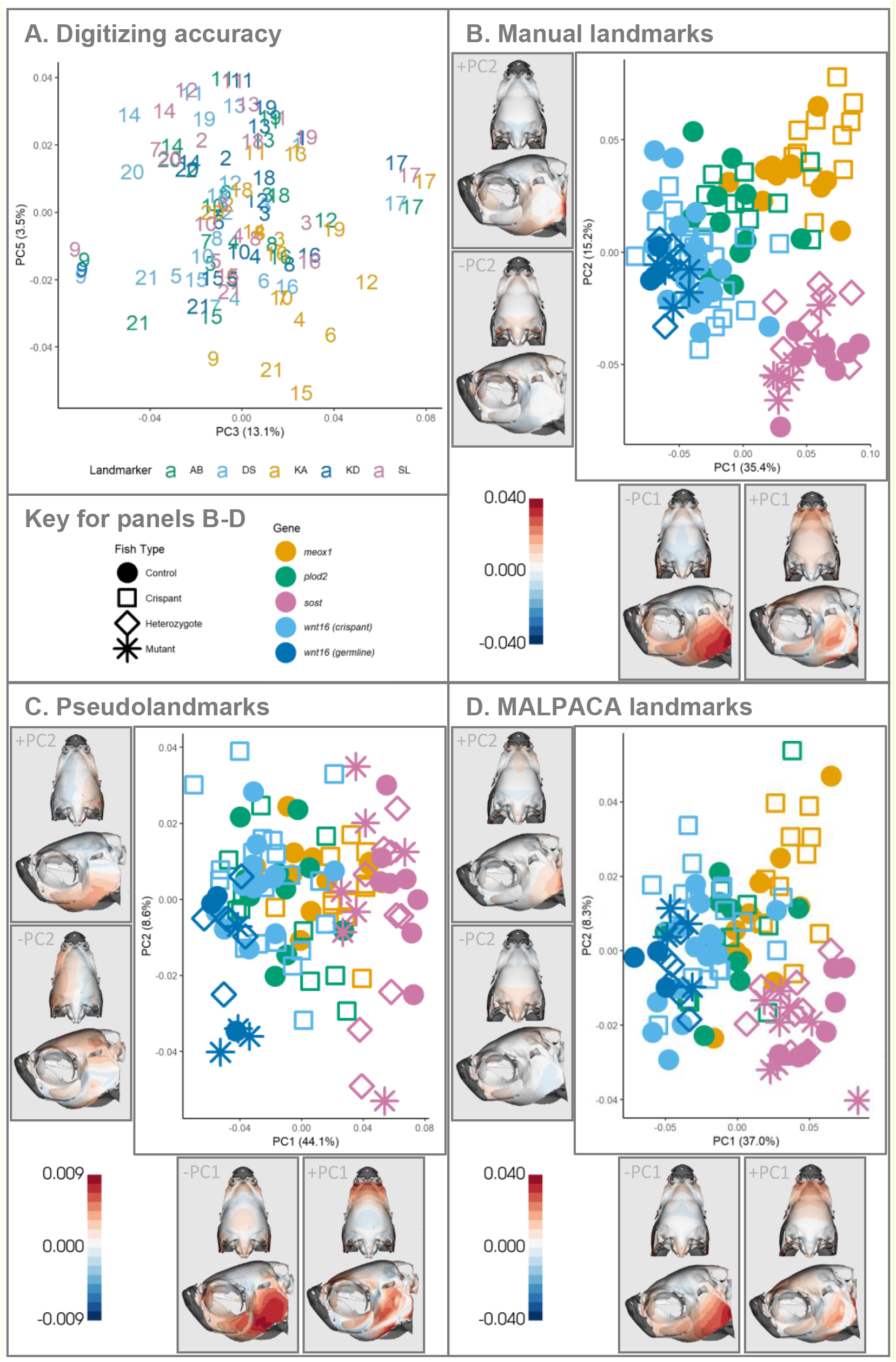
Groupwise difference plots among datasets. (A) Digitizing accuracy analysis showing PC3 and PC5 from combined manual landmarks of the *plod2* dataset, with colors indicating different individuals who landmarked datasets in this study and numbers indicating individual fish. (B-D) PC1 and PC2 from groupwise analyses using different landmarking methods including (B) 21 manually placed landmarks, 308 pseudolandmarks, and (D) 21 MALPACA placed landmarks. Point color indicates gene grouping; shape indicates type of fish. Heat maps show dorsal (top) and lateral (bottom) cranial skeletons, with colors representing Procrustes distances between mean shape and the extreme shapes across each PC. All axis labels show the percentage of variation represented by each PC.

Before examining the shape differences within any single gene, we first explored how shape varied across our complete dataset for both manual and pseudolandmarks. We made two predictions about these analyses. We predicted that control fish from the same facility (*plod2, meox1*, and *wnt16*) should be grouped closer together in morphospace as these control fish should have more similar genetic backgrounds (and all functional genes), than any control fish has to the mutant groups. Alternatively, the fish within each mutant line should be closer in morphospace to the control fish from their respective experiment, as controls are from the same clutches as the mutant.

When we run all groups, including all genes and types of fish (control, crispant, heterozygote, mutant) in a single analysis, we find that overall shape varies among groups for both the manual (F=9.658, Z=9.130, p=0.001) and pseudolandmark datasets (F=7.045, Z=8.260, p=0.001). In both analyses, groups varied across 7 of the first 9 principal components (Figures 1, S1, S2; Tables S2, S3), with the clearest patterns of divergence between genes observed in PC1 and PC2 (Figure 1B, C). Regardless of the type of fish, the first principal component of both analyses separates the *wnt16* fish on the negative side of PC1 from the *sost* and *meox1* fish on the positive side of PC1 (Figure 1B, C). This result negates our first prediction, that control fish from the same facility should be closer in morphospace, but supports our second prediction, that there is a clutch effect for craniofacial phenotype.

Across PC1 shape differences for both sets of landmarks are concentrated in the ventral portion of the opercula, and the intraocular region of the frontal bone (Figure 1B, C). Based on their positions in our morphospaces, the *wnt16* clutches (crispant, mutant, and control fish) have larger changes in the opercular region, while *sost* and *meox1* clutches have larger differences in the intraocular regions compared to the mean skull shape (Figure 1B, C). In the manual landmark dataset, PC2 separates the *sost* clutch on the negative end of the axis from all other clutches (Figure 1B). As the *sost* clutch was raised in a separate facility from all other clutches, this result provides evidence of the influence of background genetic variation and/or housing conditions on craniofacial shape. While we do not find similar separations across PC2 of the pseudolandmark analysis, we do find similar separation of clutches across PC1 (Figure 1C), and similar patterns in the MALPACA generated points (Figure 1D).

As an extra precaution to ensure these patterns are not due to different digitizers placing landmarks across the different groups of genes, we also ran MALPACA (see methods for full details). Again, we found overall shape differs between groups (F=5.828, Z=7.234, p=0.001), and similar patterns in the PCs of the MALPACA dataset compared to the manually placed landmarks (Figure 1D; Table S2).

### Somatic crispant groups

We first examined the somatic crispant fish and their control clutch mates, which includes *plod2, meox1*, and *wnt16* groups. We had larger sample sizes for these experiments than we did for germline mutant experiments because previous work on the axial skeleton found that somatic mutants have smaller effect sizes than germline mutants (Hur et al., 2017; Watson et al., 2020). The current study is more exploratory, and hence we did not have a good metric to run a similar power analysis, so we have assumed that the effect sizes of the craniofacial morphology will be similar to that of the axial skeleton. For each gene, we found differences in overall shape between control and crispant groups. Below we report the results of our pseudolandmark analyses and discuss the relationship between our results and human disease phenotypes. In general, all manual landmark analyses are aligned with the pseudolandmark analysis unless otherwise noted and details of these ground truth datasets can be found in the supplemental information (Figure S4; Table S2).

#### (A) plod2

We found strong evidence that mutations in *plod2* alter craniofacial shape. In the *plod2* pseudolandmark dataset, overall shape differed between *plod2* mutant and control fish (F=1.634, Z=1.756, p=0.041). Along PC2 control fish show patterns of variation in the posterior cranial skeleton compared to the *plod2* fish (Figure 2A; Table S3). We did not find evidence of fluctuating asymmetry among groups for *plod2* (F= 0.994, Z=0.186, p= 0.435). However, we did find differences in the symmetric component of shape variation between *plod2* and control fish (F= 2.011, Z=1.840, p= 0.039). For the symmetric component of shape variation, groups differ along the PC2 axis, again with differences concentrated in the posterior skull, especially in the opercular region (Figure 2B; Table S4).

**Figure 2.**
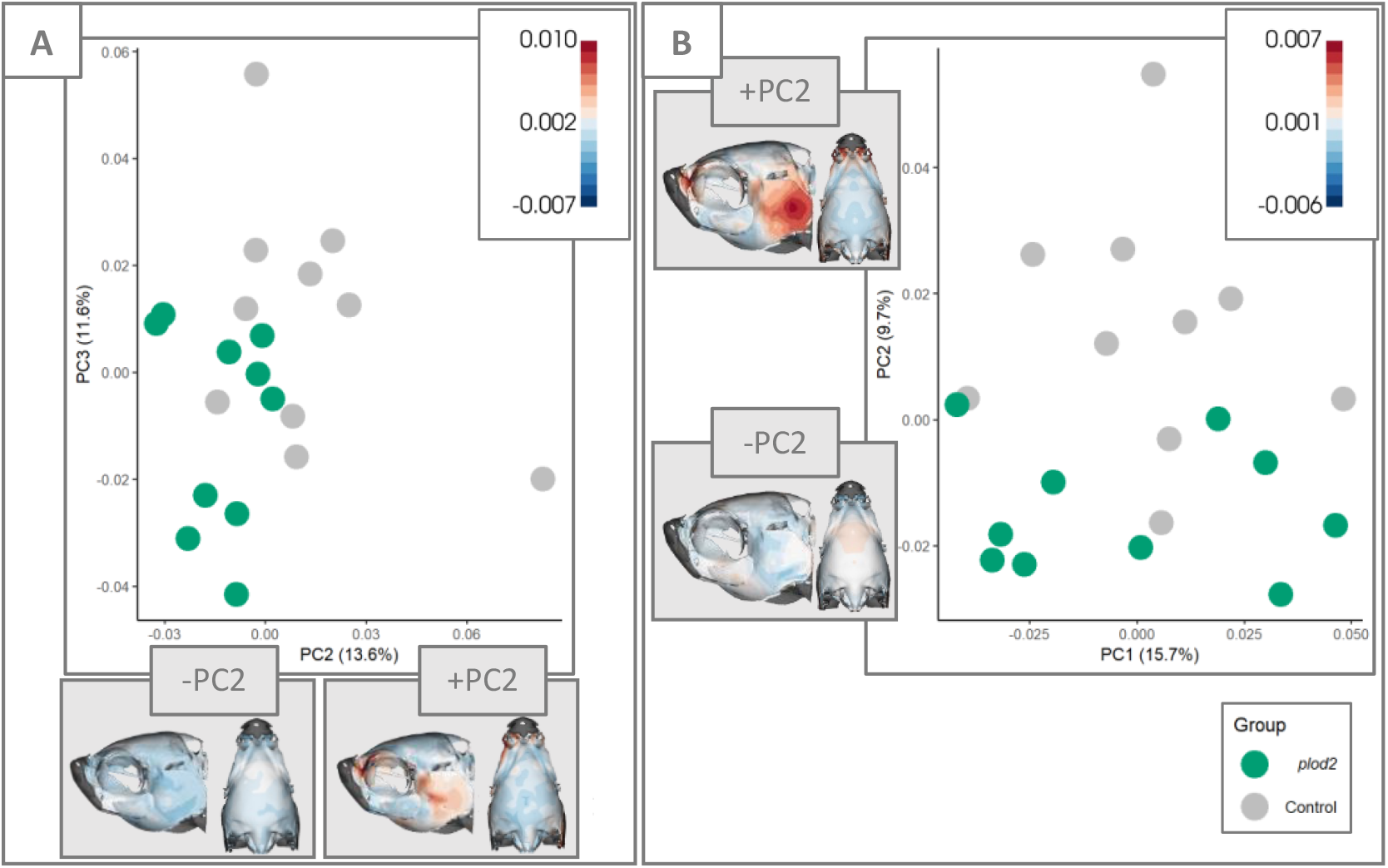
Principal component plots from analyses that show separation of *plod2* crispants and their control siblings. (A) PC2 and PC3 from pseudolandmarks, and (B) PC1 and PC2 from symmetrical components of shape variation. Point color indicates groupings. Heatmaps represent lateral (left) and dorsal (right) cranial skeletons and show Procrustes distances between mean shape and the extreme shapes across each PC. Axis labels show the percentage of variation represented by each PC.

Bruck syndrome, a rare autosomal recessive congenital form of osteogenesis imperfecta (brittle bone disease), has been associated with the human *PLOD2* gene (Lv et al., 2018). Without a functional *PLOD2* gene, bones become brittle due to the underhydroxylation effects on bone collagen telopeptide lysines during bone development (Gistelinck et al., 2021). Patients with Bruck syndrome present with large joint flexion contractures, skeletal anomalies, short stature, bone fragility, clubfoot, and joint movement restrictions (Gistelinck et al., 2016; Lv et al., 2018; Zhou et al., 2014). Unlike other forms of osteogenesis imperfecta, patients with Bruck syndrome do not present with dentinogenesis imperfecta or hearing loss (Lv et al., 2018). Considering there are fewer craniofacial symptoms in human Bruck syndrome patients, it is probable that we would not find craniofacial differences in *plod2* crispant fish. However, we chose to examine *plod2* crispants because of the clear human homologies present in the axial skeleton. Crispant zebrafish with somatic *plod2* mutations had overmineralized axial skeletons with multiple rib fracture callus, vertebral fusions, kyphosis, scoliosis, and overall decreased body length compared to control fish (Gistelinck et al., 2018; Hur et al., 2017). While we did not measure tissue mineral density in this study, we did find differences in skull shape concentrated in the opercular region (Figures 2, S4A).

#### (B) meox1

We found strong evidence that mutations in *meox1* alter craniofacial shape. Groups differed in overall shape for our 308 pseudolandmark dataset (F=2.581, Z=2.348 p=0.007), with PC1 and PC3 differing between *meox1* fish and their wildtype siblings (Figure 3A; Table S3). PC1 shows shape differences are concentrated in the subocular and intraocular portions of the skull (Figure 3A). PC3 has lower magnitudes of shape change, with shape differences in the parietal, frontal bones and ventral opercular regions (Figure 3A).

**Figure 3.**
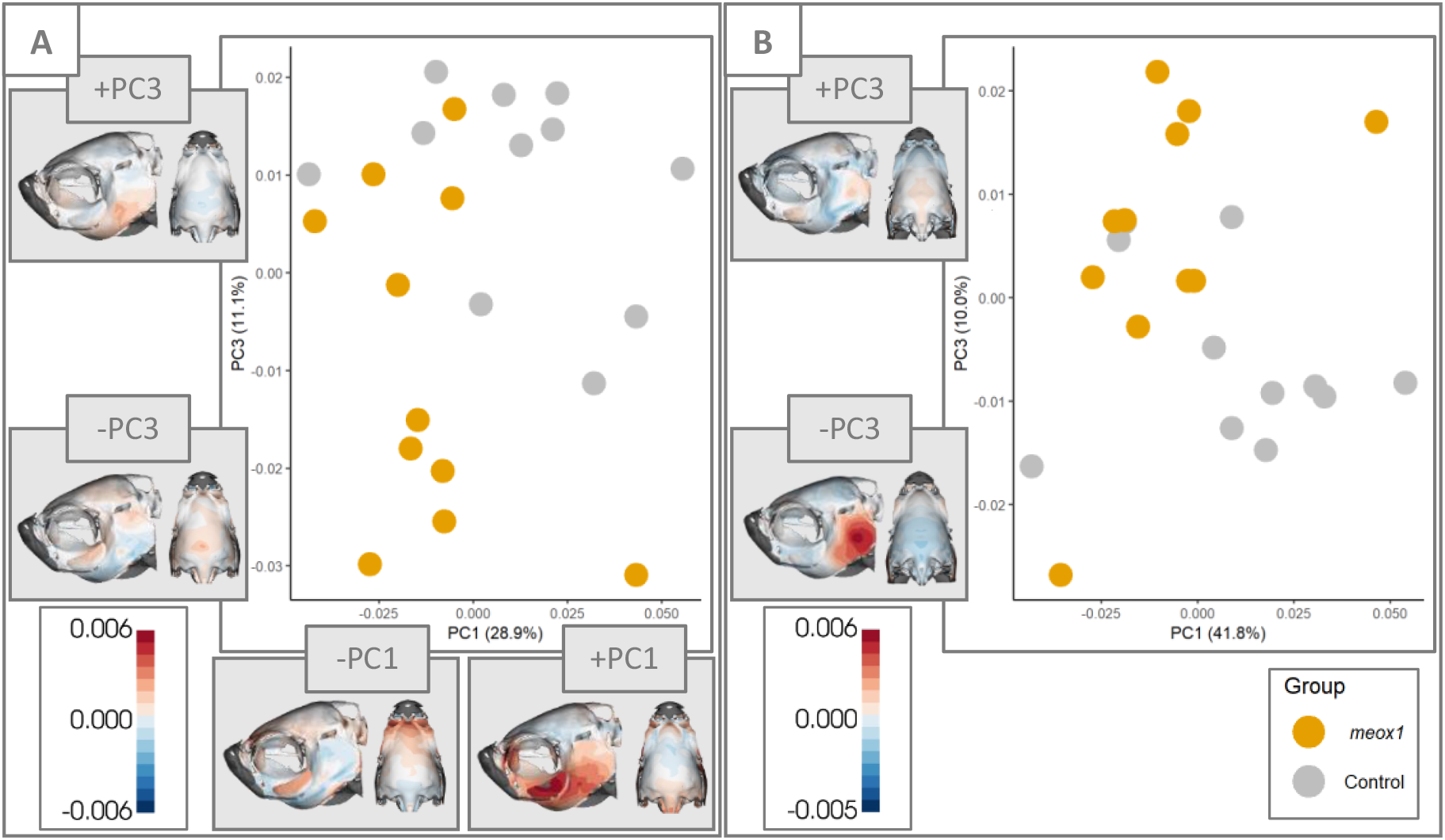
Principal component plots from analyses that show separation of *meox1* crispants and their control siblings. (A) PC1 and PC3 from pseudolandmarks, and (B) PC1 and PC3 from symmetrical components of shape variation. Point color indicates groupings. Heatmaps represent lateral (left) and dorsal (right) cranial skeletons and show Procrustes distances between mean shape and the extreme shapes across each PC. Axis labels show the percentage of variation represented by each PC.

Mesenchyme homeobox 1 is a transcription factor expressed in somites and known to be involved in the development of vertebral elements (Skuntz et al., 2009). Crispant *meox1* zebrafish have parallel symptoms to the human Klippel–Feil syndrome, a skeletal disease associated with the orthologous gene *MEOX1* (Bayrakli et al., 2013; Mohamed et al., 2013). Both human and zebrafish with nonfunctional *meox1* have symptoms that include the fusion of at least two cervical vertebrae, asymmetry in the pectoral girdle, and congenital scoliosis (Dauer et al., 2018). In contrast to humans and zebrafish, *meox1* loss-of-function mice neither develop pectoral girdle asymmetry nor exhibit congenital scoliosis (Skuntz et al., 2009), suggesting zebrafish may be a better model for Klippel-Feil Syndrome.

Given this phenotype, we anticipated an asymmetric craniofacial phenotype in the affected fish. However, the *meox1* fish do not differ from controls in fluctuating asymmetry (F=1.358, Z=0.920, p=0.185); though we did find a slight trend between groups for the symmetric component of shape variation (F=1.893, Z=1.426, p=0.09). Groups vary along the PC3 axis for our symmetry analysis, with the largest differences in shape occurring in the opercular and parietal regions (Figure 3B; Table S4). The lack of asymmetry in the skull could be attributed to differences in musculature between the axial and cranial skeletons. Nguyen and colleagues found that *meox1* controls clonal drift during muscle growth in the axial skeleton of zebrafish (Nguyen et al., 2017). Considering the differences in musculature patterns between these two skeletal systems, it is possible that *meox1* affects the axial musculoskeletal system more asymmetrically than the cranial musculoskeletal system.

#### (C) wnt16

We found little evidence that somatic mutations in *wnt16* alter craniofacial shape. Pseudolandmarks do not show any differences in overall shape between *wnt16* crispants and control fish (F=1.366, Z=1.035, p=0.152). Nor do we find differences in fluctuating asymmetry between groups (F=0.616, Z=-0.757, p=0.760). We did find a potential trend in the symmetric differences in shape variation between *wnt16* crispant and control fish (F=1.894, Z=1.439, p=0.088), with groups varying across PC2. Shape differences across PC2 are also concentrated in the occipital and subocular regions, with additional variation observed in the posterior region of the opercula (Figure 4; Table S4).

**Figure 4.**
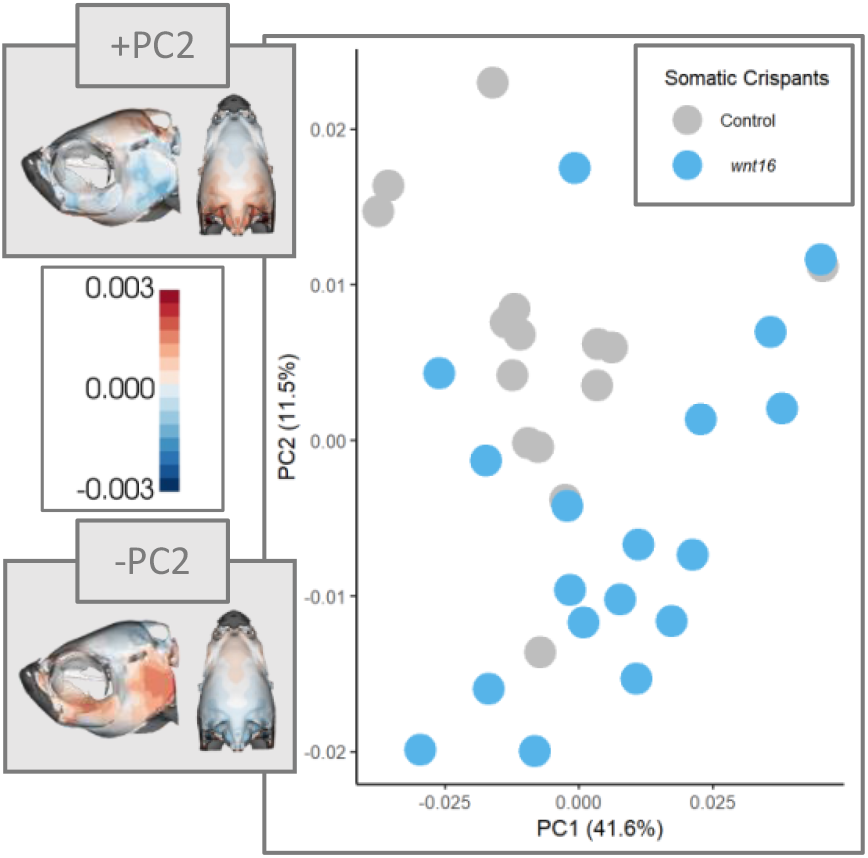
PC1 and PC2 from symmetrical components of shape variation. Point color indicates groupings. Heatmaps represent lateral (left) and dorsal (right) cranial skeletons and show Procrustes distances between mean shape and the extreme shapes across each PC. Axis labels show the percentage of variation represented by each PC.

In skeletal tissue, the Wnt signaling pathway is involved in the differentiation of osteoblasts, osteoclasts, and chondrocytes. (Ugartondo et al., 2022) The *wnt16* ligand is a regulator of bone homeostasis and has been associated with numerous skeletal phenotypes related to bone density, strength, fracture risk, and bone loss with aging (Gori et al., 2015; Movérare-skrtic et al., 2014; Todd et al., 2015; Ugartondo et al., 2022). Additionally, recent evidence from crispant zebrafish models support the potential for *wnt16* to exert pleiotropic effects on bone and lean muscle mass through a function in myogenic precursors. (Watson et al., 2021). With the exception of the occipital region, all other areas of the skull that varying between *wnt16* crispants and controls are associated with the expansion of the oral cavity required for suction feeding in fishes (Day et al., 2015). The pleotropic effect on bone and muscle, as Watson and colleagues suggest, could explain the small pattern we observe due to greater muscle use and development in the regions of the skull used for feeding (Watson et al., 2021).

### Germline mutant groups

The germline mutant fish and their control clutch mates includes *wnt16* and *sost* groups. Below we report the results of our pseudolandmark analyses and discuss the relationship between our results and human disease phenotypes.

#### (A) wnt16

Similar to our results for *wnt16* crispants, for germline mutants, we found modest evidence that mutations in *wnt16* alter craniofacial shape. The mutant grouping included *wnt16* mutants, *wnt16* heterozygotes, and controls. Past studies on zebrafish *wnt16* germline mutant axial skeletons identified phenotypes that included variable tissue mineral density, susceptibility to spontaneous fractures, and accumulation of bone calluses at an early age (McGowan et al., 2021). In the current study, we did not find strong differences in overall shape for the manual landmarks for all three groups for this set (F=1.552, Z=1.557, p=0.067), but did find differences between the mutant versus controls (F=2.102, Z=1.955, p=0.019) and mutant versus heterozygotes (F=1.976, Z=2.165, p=0.015), but not between heterozygotes versus controls (F=0.819, Z=-0.299, p=0.620). The *wnt16* mutants show the most variation in the ventral opercular and occipital regions when compared to the control or the heterozygotes (Figure 5; Table S2).

**Figure 5.**
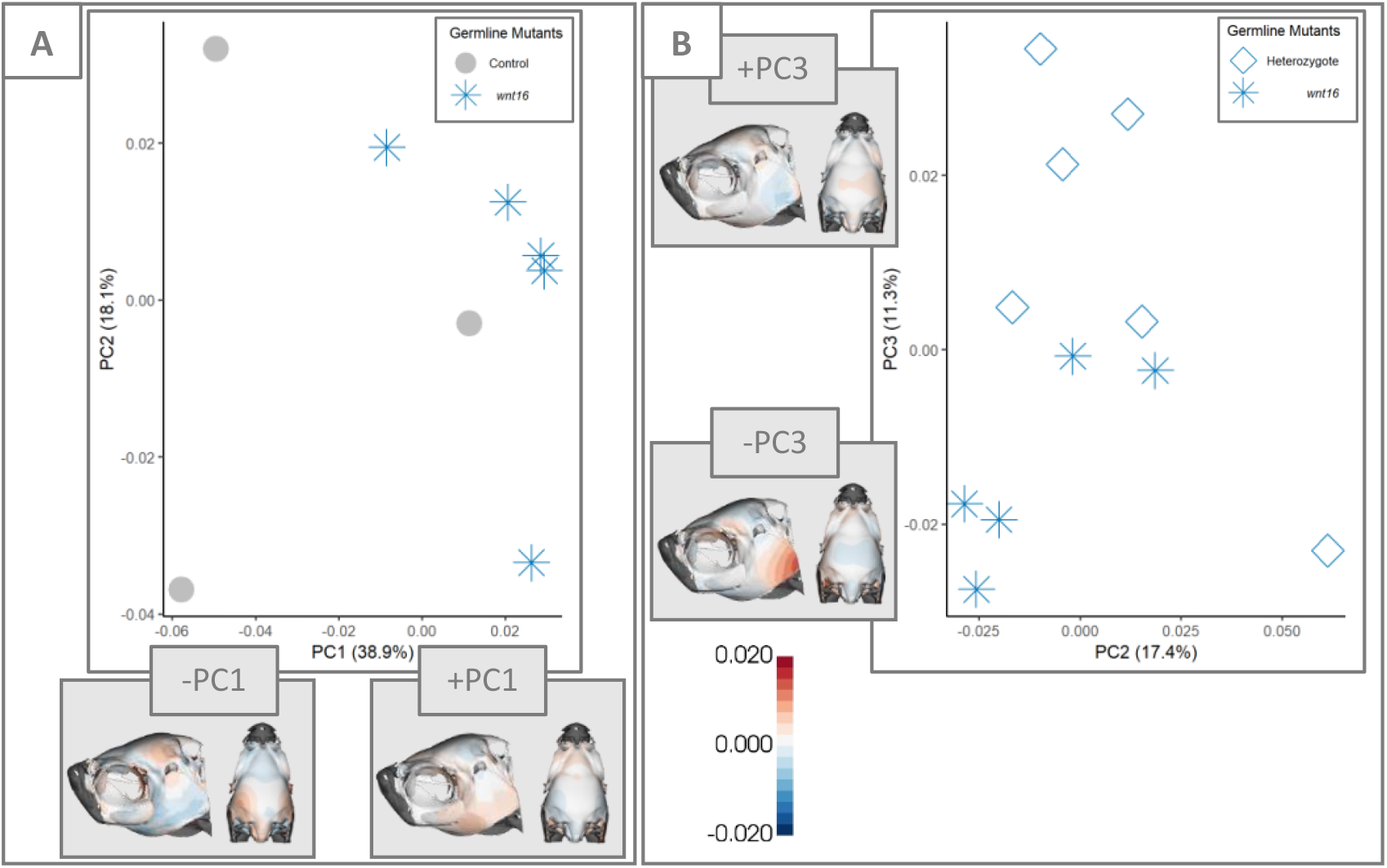
Principal component plots from manual landmark analyses that show separation of *wnt16* mutants and their (A) control and (B) heterozygote siblings. Point color and shape indicate groupings. Heatmaps represent lateral (left) and dorsal (right) cranial skeletons and show Procrustes distances between mean shape and the extreme shapes across each PC. PC. Axis labels show the percentage of variation represented by each PC.

Unlike the other three genes in this study, our pseudolandmark pipeline did not find differences in overall shape (F=0.888, Z=-0.398, p=0.655) nor asymmetric (F= 1.066, Z=0.327, p=0.380) or symmetric shape variation (F=0.971, Z=0.111, p=0.456) among the pseudolandmark points for any of the mutant groups.

#### (B) sost

We did not find strong evidence that mutations in *sost* alter craniofacial shape. Manual landmarks for *sost* mutants, heterozygotes, and control fish do not show differences in overall shape between the three groups (F=1.094, Z=0.486, p=0.314). Neither did we find differences between the three groups in overall shape when using the 308 pseudolandmarks (F=1.344, Z=0.976, p=0.172). However, we do find a trend in overall shape when only the mutant *sost* and control fish are considered (F=1.898, Z=1.437, p=0.087). Groups separate along the pseudolandmark PC2 axis, with variation concentrated in the opercular and subocular regions (Figure 6A; Table S3).

**Figure 6.**
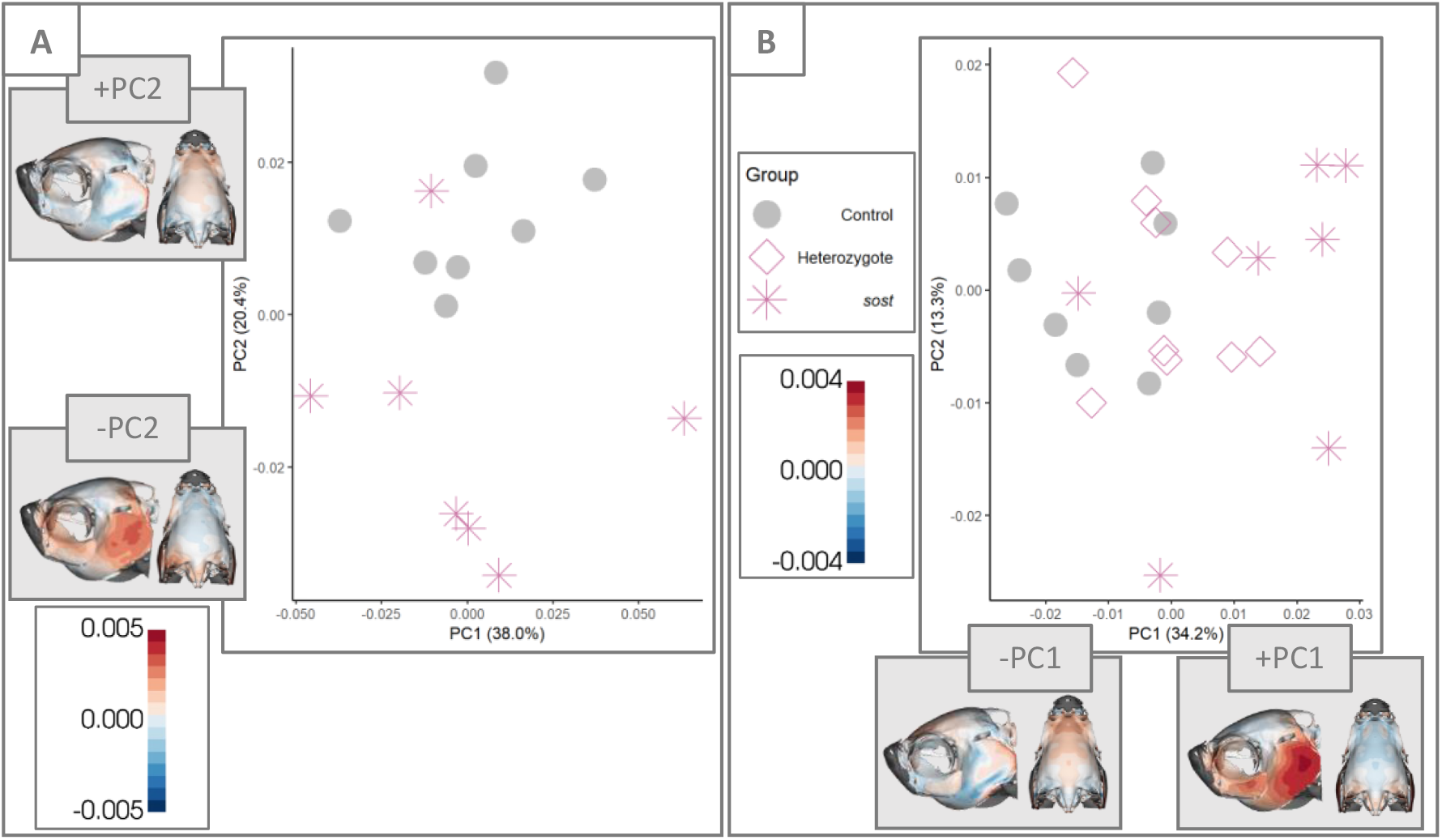
Principal component plots from manual landmark analyses that show separation of *sost* mutants, heterozygotes, and their control siblings. PC1 and PC2 from (A) pseudolandmarks and (B) symmetrical components of shape variation. Point color and shape indicates groupings. Heatmaps represent lateral (left) and dorsal (right) cranial skeletons and show Procrustes distances between mean shape and the extreme shapes across each PC. Axis labels show the percentage of variation represented by each PC.

We did not find support for differences between fluctuating asymmetry (F=0.610, Z=-0.529, p=0.677), but do find differences between groups in the symmetrical components of shape variation (F=2.272, Z=2.272, p=0.008) when all groups are considered in the same analysis, as well as when only controls and mutants are included in the analysis (F=3.624, Z=2.315, p=0.008). Groups vary across PC1, with the similar patterns as observed in the general pseudolandmark analysis (Figure 6B; Table S4). In the symmetry analysis, the heterozygous fish exhibit an intermediate phenotype between the mutant and control fish (Figure 6B).

The *SOST* gene encodes sclerostin, an osteocyte specific glycoprotein that negatively regulates bone formation through the inhibition of canonical Wnt signaling pathways in osteoblast lineage cells (Chang et al., 2014; Martínez-Gil et al., 2021). This means that in contrast to the other three genes in this study, which cause skeletal diseases as they are downregulated, *SOST* causes problems when this gene is upregulated (Kaya et al., 2022). The downregulation of *sost*/sclerostin in osteocytes has a similar association with the mechanotransduction cascade as *wnt16* upregulation because the downregulation of sclerostin activates Wnt signaling and directs osteogenesis to where bone is structurally needed (Tu et al., 2012). In mice long bones (Robling et al., 2008) and jaws (Inoue et al., 2019), the modulation of sclerostin levels appears to be a finely tuned mechanism by which osteocytes coordinate regional and local osteogenesis in response to increased mechanical stimulation. Sclerostin was first recognized in the study of two rare high bone mass disorders, sclerosteosis (Balemans et al., 2001; Brunkow et al., 2001) and Van Buchem’s disease (Van Buchem et al., 1662), but is involved in the pathogenesis of many skeletal disorders. Modulation of this osteocyte-derived negative signal is therapeutically relevant for disorders associated with bone loss (Winkler et al., 2003; Winkler et al., 2004). Due to the human disease phenotype associated with sclerostin being particularly prominent in the skull, jaw, and long bones (Sebastian and Loots, 2018), we predicted that mutant zebrafish would exhibit larger craniofacial skeletons than the controls. Our results do not support this prediction as we did not find any strong evidence that *sost* mutations alter craniofacial morphology. Mouse *sost* mutants do not have a clear craniofacial phenotype either (Sebastian and Loots, 2018), suggesting that the cranial hyperostosis phenotype may be specific to humans.

## Conclusions

In this study we tested a previously developed semiautomated pipeline (Diamond et al., 2022) to identify areas of craniofacial variation in four zebrafish models of skeletal disease. Our results suggest that regardless of the type of fish used in a study (somatic vs germline), there is a strong clutch effect in zebrafish. Clutch effects have also been observed in zebrafish cardiac performance (Schwerte et al., 2005) and behavioral phenotypes (Joo et al., 2021). Further, the *sost* fish, which were from a different facility show the greatest separation in morphospace from the other groups in our combined analysis (Figure 1B-D). Together these results suggest that non-sibling fish, or fish from different facilities should not be used as controls in quantitative phenotyping studies.

The four zebrafish models tested in this study were chosen based on previous work finding phenotypes in the axial skeleton that were homologous to known human diseases. Due to the complexity of the craniofacial skeleton, previous studies had not assessed if these models possessed a craniofacial phenotype. In two of the four model systems (*plod2* and *meox1*) our pseudolandmark pipeline identified areas of variation in the skull. In contrast to manual landmarks, which are time intensive to collect and can introduce interobserver error into the dataset (Figure 1A), our results supports that our previously established pseudolandmark pipeline allows for more efficient data collection, higher consistency and repeatability, and denser morphometric analysis (Diamond et al., 2022).

In both of the *plod2* and *meox1* models, we found the greatest variation in the opercular region of the skull. While zebrafish skulls possess both cellular and acellular bone, the opercula are neural crest-derived compact cellular bone, which is the same developmental pathway and ossification method as human craniofacial bones (Galea et al., 2021; Kague et al., 2012; Weigele and Franz-Odendaal, 2016). Additionally, the phenotypes we identified reflect the human disease phenotypes associated with these genes. A more recent GWAS study that focused on skull bone mineral density in humans has identified four novel loci that may be associated with intramembranous ossification (Medina-Gomez et al., 2021). Crispant zebrafish models for these genes would be an interesting future test of this pipeline.

Documenting and describing phenotypic variation is critical for both the evolutionary and clinical understanding of complex traits (Conith et al., 2019). By examining phenotypic differences within and among these different models, we show the effectiveness of this pipeline for broad use in phenotyping the craniofacial skeleton of zebrafish, a heavily used model in studies of development and evolutionary biology.

## Acknowledgements

We thank members of the Maga lab and MSBL for feedback and development of this project. This project was partly supported by National Science Foundation grants An Integrated Platform for Retrieval, Visualization and Analysis of 3D Morphology from Digital Biological Collections (DBI/1759883) and Biology Guided Neural Networks for discovering phenotypic traits (OAC/1939505) to AMM. RYK was supported by NIH Grant AR074417.

## Author Contributions

Contribution of fish and microCT scans: RYK. Conceptualization and methodology: all authors, manual landmark data collection: AB, DS, KA, KMD, SLL. Writing: AB, DS, KA, KMD, SLL; Editing and approval: all authors.

## Data availability

Atlas and pseudo-landmark points are available at: https://github.com/SlicerMorph/ZF_Skull_atlas/

## Literature Cited

Adams, D. C. and Otárola-Castillo, E. (2013). Geomorph: An r package for the collection and analysis of geometric morphometric shape data. Methods Ecol. Evol. 4, 393–399.

Balemans, W., Ebeling, M., Patel, N., Hul, E. Van, Olson, P., Dioszegi, M., Lacza, C., Wuyts, W., Ende, J. Van Den, Willems, P., et al. (2001). Increased bone density in sclerosteosis is due to the deficiency of a novel secreted protein (SOST). Hum. Mol. Genet. 10, 537–543.

Bayrakli, F., Guclu, B., Yakicier, C., Balaban, H., Kartal, U., Erguner, B., Sagiroglu, M. S., Yuksel, S., Ozturk, A. R., Kazanci, B., et al. (2013). Mutation in MEOX1 gene causes a recessive Klippel-Feil syndrome subtype. BMC Genet. 14, 1–7.

Brunkow, M. E., Gardner, J. C., Van Ness, J., Paeper, B. W., Kovacevich, B. R., Proll, S., Skonier, J. E., Zhao, L., Sabo, P. J., Fu, Y. H., et al. (2001). Bone dysplasia sclerosteosis results from loss of the SOST gene product, a novel cystine knot-containing protein. Am. J. Hum. Genet. 68, 577–589.

Busse, B., Galloway, J. L., Gray, R. S., Harris, M. P. and Kwon, R. Y. (2020). Zebrafish: An Emerging Model for Orthopedic Research. J. Orthop. Res. 38, 925–936.

Chang, M. K., Kramer, I., Keller, H., Gooi, J. H., Collett, C., Jenkins, D., Ettenberg, S. A., Cong, F., Halleux, C. and Kneissel, M. (2014). Reversing LRP5-dependent osteoporosis and SOST deficiency–induced sclerosing bone disorders by altering WNT signaling activity. J. Bone Miner. Res. 29, 29–42.

Conith, A. J., Lam, D. T. and Albertson, C. (2019). Muscle-induced loading as an important source of variation in craniofacial skeletal shape. Genesis 57, e23263.

Dauer, M. V. P., Currie, P. D. and Berger, J. (2018). Skeletal malformations of Meox1-deficient zebrafish resemble human Klippel–Feil syndrome. J. Anat. 233, 687–695.

Day, S. W., Higham, T. E., Holzman, R. and Van Wassenbergh, S. (2015). Morphology, Kinematics, and Dynamics: The Mechanics of Suction Feeding in Fishes. Integr. Comp. Biol. 55, 21–35.

Diamond, K. M., Rolfe, S. M., Kwon, R. Y. and Maga, A. M. (2022). Computational anatomy and geometric shape analysis enables analysis of complex craniofacial phenotypes in zebrafish. Biol. Open 11, bio058948.

Dietrich, K., Fiedler, I. A. K., Kurzyukova, A., López-Delgado, A. C., McGowan, L. M., Geurtzen, K., Hammond, C. L., Busse, B. and Knopf, F. (2021). Skeletal biology and disease modeling in zebrafish. J. Bone Miner. Res. 36, 436–458.

Fedorov, A., Beichel, R., Kalpathy-Cramer, J., Finet, J., Jc, F.-R., Pujol, S., Bauer, C., Jennings, D., Fennessy, F., Sonka, M., et al. (2012). 3D Slicer as an image computing platform for the quantitative imaging network. Magn Reson Imaging 30, 1323–1241.

Galea, G. L., Zein, M. R., Allen, S. and Francis-West, P. (2021). Making and shaping endochondral and intramembranous bones. Dev. Dyn. 250, 414–449.

Garg, B., Tomar, N., Biswas, A., Mehta, N. and Malhotra, R. (2022). Understanding Musculoskeletal Disorders Through Next-Generation Sequencing. J. Bone Jt. Surg. 10, e21.00165.

Gistelinck, C., Witten, P. E., Huysseune, A., Symoens, S., Malfait, F., Larionova, D., Simoens, P., Dierick, M., Van Hoorebeke, L., De Paepe, A., et al. (2016). Loss of Type I Collagen Telopeptide Lysyl Hydroxylation Causes Musculoskeletal Abnormalities in a Zebrafish Model of Bruck Syndrome. J. Bone Miner. Res. 31, 1930–1942.

Gistelinck, C., Kwon, R. Y., Malfait, F., Symoens, S., Harris, M. P., Henke, K., Hawkins, M. B., Fisher, S., Sips, P., Guillemyn, B., et al. (2018). Zebrafish type I collagen mutants faithfully recapitulate human type I collagenopathies. Proc. Natl. Acad. Sci. U. S. A. 115, E8037–E8046.

Gistelinck, C., Weis, M. A., Rai, J., Schwarze, U., Niyazov, D., Song, K. M., Byers, P. H. and Eyre, D. R. (2021). Abnormal bone collagen cross-linking in Osteogenesis Imperfecta/Bruck Syndrome caused by compound heterozygous PLOD2 mutations. JBMR Plus 5, 1–15.

Goodall, C. (1991). Procrustes methods in the statistical analysis of shape. J. R. Stat. Soc. 53, 285–339.

Gori, F., Lerner, U., Ohlsson, C. and Baron, R. (2015). A new WNT on the bone: WNT16, cortical bone thickness, porosity and fractures. Bonekey Rep. 4, 1–6.

Hamersma, H., Gardner, J. and Beighton, P. (2003). The natural history of sclerosteosis. Clin. Genet. 63, 192–197.

Hur, M., Gistelinck, C. A., Huber, P., Lee, J., Thompson, M. H., Monstad-Rios, A. T., Watson, C. J., McMenamin, S. K., Willaert, A., Parichy, D. M., et al. (2017). MicroCT-Based Phenomics in the Zebrafish Skeleton Reveals Virtues of Deep Phenotyping in a Distributed Organ System. Elife 6, e26014.

Inoue, M., Ono, T., Kameo, Y., Sasaki, F., Ono, T., Adachi, T. and Nakashima, T. (2019). Forceful mastication activates osteocytes and builds a stout jawbone. Sci. Rep. 9, 1–12.

Joo, W., Vivian, M. D., Graham, B. J., Soucy, E. R. and Thyme, S. B. (2021). A Customizable Low-Cost System for Massively Parallel Zebrafish Behavioral Phenotyping. Front. Behav. Neurosci. 14, 1–11.

Kague, E. and Karasik, D. (2022). Functional validation of osteoporosis genetic findings using small fish models. Genes (Basel). 13,.

Kague, E., Gallagher, M., Burke, S., Parsons, M., Franz-Odendaal, T. and Fisher, S. (2012). Skeletogenic Fate of Zebrafish Cranial and Trunk Neural Crest. PLoS One 7, 1–13.

Kaya, S., Schurman, C. A., Dole, N. S., Evans, D. S. and Alliston, T. (2022). Prioritization of Genes Relevant to Bone Fragility Through the Unbiased Integration of Aging Mouse Bone Transcriptomics and Human GWAS Analyses. J. Bone Miner. Res. 37, 804–817.

Kwon, R. Y., Watson, C. J. and Karasik, D. (2019). Using zebrafish to study skeletal genomics. Bone 126, 37–50.

Lv, F., Xu, X., Song, Y., Li, L., Asan, Wang J., Yang, H., Wang, O., Jiang, Y., Xia, W., et al. (2018). Novel Mutations in PLOD2 Cause Rare Bruck Syndrome. Calcif. Tissue Int. 102, 296–309.

Martínez-Gil, N., Roca-Ayats, N., Cozar, M., Garcia-Giralt, N., Ovejero, D., Nogués, X., Grinberg, D. and Balcells, S. (2021). Genetics and genomics of SOST: Functional analysis of variants and genomic regulation in osteoblasts. Int. J. Mol. Sci. 22, 1–14.

McGowan, L. M., Kague, E., Vorster, A., Newham, E., Cross, S. and Hammond, C. L. (2021). Wnt16 Elicits a Protective Effect Against Fractures and Supports Bone Repair in Zebrafish. JBMR Plus 5, 1–14.

Medina-Gomez, C., Kemp, J. P., Estrada, K., Eriksson, J., Liu, J., Reppe, S., Evans, D. M., Heppe, D. H. M., Vandenput, L., Herrera, L., et al. (2012). Meta-analysis of genome-wide scans for total body BMD in children and adults reveals allelic heterogeneity and age-specific effects at the WNT16 locus. PLoS Genet. 8,.

Medina-Gomez, C., Mullin, B. H., Chesi, A., Prijatelj, V. and John, P. (2021). Genome Wide Association Metanalysis of Skull Bone Mineral Density Identifies Loci Relevant for Osteoporosis and Craniosynostosis. medRxiv 1–38.

Mohamed, J. Y., Faqeih, E., Alsiddiky, A., Alshammari, M. J., Ibrahim, N. A. and Alkuraya, F. S. (2013). Mutations in MEOX1, encoding mesenchyme homeobox 1, cause klippel-feil anomaly. Am. J. Hum. Genet. 92, 157–161.

Mortier, G. R., Cohn, D. H., Cormier-Daire, V., Hall, C., Krakow, D., Mundlos, S., Nishimura, G., Robertson, S., Sangiorgi, L., Savarirayan, R., et al. (2019). Nosology and classification of genetic skeletal disorders: 2019 revision. Am. J. Med. Genet. Part A 179, 2393–2419.

Movérare-skrtic, S., Henning, P., Liu, X., Nagano, K., Engdahl, C., Koskela, A., Zhang, F. and Eriksson, E. E. (2014). Osteoblast-derived WNT16 represses osteoclastogenesis and prevents cortical bone fragility fractures. Nat. Med. 20, 1279–1288.

Nguyen, P. D., Gurevich, D. B., Sonntag, C., Hersey, L., Alaei, S., Nim, H. T., Siegel, A., Hall, T. E., Rossello, F. J., Boyd, S. E., et al. (2017). Muscle Stem Cells Undergo Extensive Clonal Drift during Tissue Growth via Meox1-Mediated Induction of G2 Cell-Cycle Arrest. Cell Stem Cell 21, 107-119.e6.

Porto, A., Rolfe, S. M. and Maga, A. M. (2021). ALPACA: A fast and accurate computer vision approach for automated landmarking of three-dimensional biological structures. Methods Ecol. Evol. 12, 2129–2144.

Robling, A. G., Niziolek, P. J., Baldridge, L. A., Condon, K. W., Allen, M. R., Alam, I., Mantila, S. M., Gluhak-Heinrich, J., Bellido, T. M., Harris, S. E., et al. (2008). Mechanical stimulation of bone in vivo reduces osteocyte expression of Sost/sclerostin. J. Biol. Chem. 283, 5866–5875.

Rolfe, S., Pieper, S., Porto, A., Diamond, K., Winchester, J., Shan, S., Kirveslahti, H., Boyer, D., Summers, A. and Maga, A. M. (2021). SlicerMorph: An open and extensible platform to retrieve, visualize and analyze 3D morphology. Methods Ecol. Evol. 12, 1816–1825.

Schlager, S. (2017). Morpho and Rvcg - Shape Analysis in R: R-Packages for Geometric Morphometrics, Shape Analysis and Surface Manipulations. In Statistical Shape and Deformation Analysis: Methods, Implementation and Applications, pp. 217–256. Elsevier Ltd.

Schwerte, T., Voigt, S. and Pelster, B. (2005). Epigenetic variations in early cardiovascular performance and hematopoiesis can be explained by maternal and clutch effects in developing zebrafish (Danio rerio). Comp. Biochem. Physiol. A 141, 200–209.

Sebastian, A. and Loots, G. G. (2018). Genetics of Sost/SOST in sclerosteosis and van Buchem disease animal models. Metabolism. 80, 38–47.

Skuntz, S., Mankoo, B., Nguyen, M. T. T., Hustert, E., Nakayama, A., Tournier-Lasserve, E., Wright, C. V. E., Pachnis, V., Bharti, K. and Arnheiter, H. (2009). Lack of the mesodermal homeodomain protein MEOX1 disrupts sclerotome polarity and leads to a remodeling of the cranio-cervical joints of the axial skeleton. Dev. Biol. 332, 383–395.

Thomas, E. D. and Raible, D. W. (2019). Distinct progenitor populations mediate regeneration in the zebrafish lateral line. Elife 8, 1–27.

Todd, H., Galea, G. L., Meakin, L. B., Delisser, P. J., Lanyon, L. E., Windahl, S. H. and Price, J. S. (2015). Wnt16 is associated with age-related bone loss and estrogen withdrawal in murine bone. PLoS One 10, 1–16.

Tu, X., Rhee, Y., Condon, K., Bivi, N., Allen, M. R., Stolina, M., Turner, C. H., Robling, A. G., Plotkin, L. I. and Bellido, T. (2012). Sost downregulation and local Wnt signaling are required for the osteogenic response to mechanical loading. Bone 50, 209–217.

Ugartondo, N., Grinberg, D. and Balcells, S. (2022). Wnt Pathway Extracellular Components and Their Essential. 1, 1–52.

Van Buchem, F. S. P., Hadders, H. N., Hansen, J. F. and Woldring, M. G. (1662). Hyperostosis corticalis generalisata: report of seven cases. Am. J. Med. 33, 387–397.

Watson, C. J., Monstad-Rios, A. T., Bhimani, R. M., Gistelinck, C., Willaert, A., Coucke, P., Hsu, Y. H. and Kwon, R. Y. (2020). Phenomics-Based Quantification of CRISPR-Induced Mosaicism in Zebrafish. Cell Syst. 10, 275-286.e5.

Watson, C. J., Morphin Montes de Oca, E., Fiedler, I. A. K., Cronrath, A. R., Callies, L. K., Swearer, A. A., Sethuraman, V., Ahmed, A. R., Monstad-Rios, A. T., Rojas, M. F., et al. (2021). wnt16 exerts pleiotropic effects on bone and lean mass in zebrafish. bioRxiv doi:10.1101/2021.08.12.456120.

Weigele, J. and Franz-Odendaal, T. A. (2016). Functional bone histology of zebrafish reveals two types of endochondral ossification, different types of osteoblast clusters and a new bone type. J. Anat. 229, 92–103.

Winkler, D. G., Sutherland, M. K., Geoghegan, J. C., Yu, C., Hayes, T., Skonier, J. E., Shpektor, D., Jonas, M., Kovacevich, B. R., Staehling-Hampton, K., et al. (2003). Osteocyte control of bone formation via sclerostin, a novel BMP antagonist. EMBO J. 22, 6267–6276.

Winkler, D. G., Yu, C., Geoghegan, J. C., Ojala, E. W., Skonier, J. E., Shpektor, D., Sutherland, M. K. and Latham, J. A. (2004). Noggin and sclerostin bone morphogenetic protein antagonists form a mutually inhibitory complex. J. Biol. Chem. 279, 36293–36298.

Zhang, C., Porto, A., Rolfe, S., Kocatulum, A. and Maga, A. M. (2022). Automated Landmarking via Multiple Templates. bioRxiv 2022.01.04.474967.

Zhou, P., Liu, Y., Lv, F., Nie, M., Jiang, Y., Wang, O., Xia, W., Xing, X. and Li, M. (2014). Novel mutations in FKBP10 and PLOD2 cause rare bruck syndrome in Chinese Patients. PLoS One 9, 1–8.

