## Supplemental Information for "Examining craniofacial variation among crispant and mutant zebrafish models of human skeletal diseases"

**Table S1.** 21 manual landmark points. Images of these landmark points can be found in (Diamond et al., 2022).

| Landmark | Definition |
|---|---|
| 1 | Anterior frontal |
| 2 | Dorsal epiphyseal bar |
| 3 | Dorsal meeting of frontal/parietal |
| 4 | Left meeting of parietal/supraoccipital |
| 5 | Dorsal meeting of parietal/supraoccipital |
| 6 | Right meeting of parietal/supraoccipital |
| 7 | Dorsal cranium - 1st vertebrae |
| 8 | Left epiphyseal bar |
| 9 | Left meeting of frontal/parietal |
| 10 | Left posterior pterotic |
| 11 | Left meeting of opercle/hyomandibular |
| 12 | Left meeting of opercle/interopercle |
| 13 | Left posterior opercula |
| 14 | Left meeting of dentary/quadrate |
| 15 | Right epiphyseal bar |
| 16 | Right meeting of frontal/parietal |
| 17 | Right posterior pterotic |
| 18 | Right meeting of opercle/hyomandibular |
| 19 | Right meeting of opercle/interopercle |
| 20 | Right posterior opercula |
| 21 | Right meeting of dentary/quadrate |

**Table S2.** Procrustes ANOVA results for analyses of 21 manually landmarked datasets in this study. The first grouping is for test if PCs vary by the individual who landmarked the *plod2* dataset (PC ~ individual). The remaining groupings test if shape coordinates and principal components differ between, crispant/ mutant and controls (Component ~ group). The proportion of variance explained by each PC is given in the %Var column.

| PC | % Var | DF | F | Z | p |
|---|---|---|---|---|---|
| **Digitizing Accuracy of *plod2*** | | | | | |
| PC1 | 19.1 | 4 | 0.075 | -2.332 | 0.992 |
| PC2 | 16.7 | 4 | 1.152 | 0.412 | 0.341 |
| PC3 | 13.1 | 4 | 3.181 | 2.144 | 0.016 |
| PC4 | 9.6 | 4 | 0.136 | -1.951 | 0.975 |
| PC5 | 6.9 | 4 | 3.204 | 2.115 | 0.012 |
| PC6 | 4.7 | 4 | 0.505 | -0.603 | 0.724 |
| PC7 | 3.5 | 4 | 1.638 | 0.998 | 0.154 |
| PC8 | 3.2 | 4 | 0.781 | -0.141 | 0.555 |
| PC9 | 3.1 | 4 | 3.270 | 2.065 | 0.022 |
| PC10 | 2.6 | 4 | 1.183 | 0.462 | 0.326 |
| PC11 | 2.4 | 4 | 1.923 | 1.205 | 0.120 |
| **all groups** | | | | | |
| PC1 | 35.4 | 11 | 42.346 | 15.086 | 0.001 |
| PC2 | 15.2 | 11 | 29.316 | 10.643 | 0.001 |
| PC3 | 8.7 | 11 | 2.837 | 2.760 | 0.004 |
| PC4 | 5.4 | 11 | 4.874 | 4.202 | 0.001 |
| PC5 | 5.0 | 11 | 2.011 | 1.811 | 0.037 |
| PC6 | 4.4 | 11 | 3.563 | 3.337 | 0.001 |
| PC7 | 3.0 | 11 | 6.737 | 5.146 | 0.001 |
| PC8 | 2.2 | 11 | 0.791 | -0.352 | 0.627 |
| **MALPACA** | | | | | |
| PC1 | 37.0 | 11 | 23.046 | 8.794 | 0.001 |
| PC2 | 8.3 | 11 | 6.954 | 5.850 | 0.001 |
| PC3 | 8.6 | 11 | 2.992 | 2.991 | 0.003 |
| PC4 | 5.0 | 11 | 2.052 | 1.871 | 0.033 |
| PC5 | 4.5 | 11 | 3.114 | 2.881 | 0.002 |
| PC6 | 4.4 | 11 | 2.515 | 2.326 | 0.007 |
| PC7 | 3.1 | 11 | 2.333 | 2.119 | 0.019 |
| PC8 | 2.6 | 11 | 2.701 | 2.658 | 0.006 |
| PC9 | 2.5 | 11 | 1.227 | 0.545 | 0.294 |
| PC10 | 2.1 | 11 | 0.373 | -1.779 | 0.967 |
| ***plod2* crispant vs control** | | | | | |
| PC1 | 23.6 | 1 | 4.916 | 1.712 | 0.031 |
| PC2 | 19.1 | 1 | 6.139 | 1.948 | 0.023 |
| PC3 | 12.8 | 1 | 0.203 | -0.403 | 0.653 |
| PC4 | 11.4 | 1 | 0.000 | -2.282 | 0.993 |

| | | | | | |
|---|---|---|---|---|---|
| PC5 | 6.4 | 1 | 4.872 | 1.633 | 0.045 |
| PC6 | 5.0 | 1 | 0.341 | -0.145 | 0.565 |
| PC7 | 3.9 | 1 | 0.070 | -0.862 | 0.798 |
| PC8 | 3.3 | 1 | 0.000 | -2.268 | 0.986 |
| PC9 | 3.0 | 1 | 1.426 | 0.707 | 0.254 |
| PC10 | 2.3 | 1 | 0.197 | -0.466 | 0.681 |
| ***meox1* crispant vs control** | | | | | |
| PC1 | 24.4 | 1 | 11.428 | 2.533 | 0.001 |
| PC2 | 15.1 | 1 | 0.037 | -1.114 | 0.858 |
| PC3 | 13.5 | 1 | 0.076 | -0.749 | 0.753 |
| PC4 | 10.0 | 1 | 0.146 | -0.503 | 0.682 |
| PC5 | 6.9 | 1 | 1.244 | 0.626 | 0.282 |
| PC6 | 6.0 | 1 | 3.206 | 1.382 | 0.082 |
| PC7 | 4.3 | 1 | 0.389 | -0.080 | 0.533 |
| PC8 | 3.9 | 1 | 0.021 | -1.305 | 0.896 |
| PC9 | 3.5 | 1 | 2.050 | 0.900 | 0.196 |
| PC10 | 2.9 | 1 | 2.830 | 1.276 | 0.096 |
| PC11 | 2.1 | 1 | 0.223 | -0.372 | 0.646 |
| ***wnt16* crispant vs control** | | | | | |
| PC1 | 30.1 | 1 | 0.043 | -1.083 | 0.845 |
| PC2 | 15.1 | 1 | 6.406 | 1.976 | 0.015 |
| PC3 | 11.8 | 1 | 0.080 | -0.843 | 0.797 |
| PC4 | 7.5 | 1 | 2.952 | 1.298 | 0.090 |
| PC5 | 4.9 | 1 | 0.515 | 0.144 | 0.452 |
| PC6 | 4.0 | 1 | 1.684 | 0.861 | 0.203 |
| PC7 | 3.5 | 1 | 2.688 | 1.269 | 0.114 |
| PC8 | 2.9 | 1 | 3.933 | 1.572 | 0.053 |
| PC9 | 2.4 | 1 | 2.912 | 1.333 | 0.092 |
| PC10 | 2.1 | 1 | 0.227 | -0.386 | 0.656 |
| PC11 | 2.1 | 1 | 4.558 | 1.721 | 0.036 |
| ***wnt16* mutant vs control** | | | | | |
| PC1 | 38.9 | 1 | 7.648 | 1.694 | 0.021 |
| PC2 | 18.1 | 1 | 0.048 | -0.982 | 0.834 |
| PC3 | 14.9 | 1 | 0.001 | -2.289 | 0.995 |
| PC4 | 9.9 | 1 | 3.623 | 1.278 | 0.121 |
| PC5 | 7.4 | 1 | 0.023 | -1.503 | 0.940 |
| PC6 | 6.5 | 1 | 0.090 | -1.023 | 0.847 |
| PC7 | 4.3 | 1 | 0.228 | -0.308 | 0.635 |
| ***wnt16* mutant vs heterozygote** | | | | | |
| PC1 | 42.9 | 1 | 2.495 | 1.465 | 0.069 |
| PC2 | 17.4 | 1 | 1.967 | 0.948 | 0.170 |
| PC3 | 11.3 | 1 | 5.607 | 1.561 | 0.044 |
| PC4 | 7.7 | 1 | 1.117 | 0.468 | 0.324 |
| PC5 | 5.6 | 1 | 0.100 | -0.623 | 0.736 |

| | | | | | |
|---|---|---|---|---|---|
| PC6 | 5.3 | 1 | 0.023 | -1.383 | 0.920 |
| PC7 | 3.4 | 1 | 0.240 | -0.290 | 0.618 |
| PC8 | 2.8 | 1 | 0.606 | 0.121 | 0.447 |

**Table S3.** Procrustes ANOVA results for analyses of 308 pseudolandmark datasets in this study. The groupings test if PCs differ between crispant/mutant and controls (PC ~ group). The proportion of variance explained by each PC is given in the %Var column.

| PC | % Var | DF | F | Z | p |
|---|---|---|---|---|---|
| **all groups** | | | | | |
| PC1 | 44.1 | 11 | 29.922 | 11.826 | 0.001 |
| PC2 | 8.6 | 11 | 2.051 | 1.913 | 0.033 |
| PC3 | 5.9 | 11 | 2.765 | 2.697 | 0.004 |
| PC4 | 4.7 | 11 | 2.475 | 2.441 | 0.009 |
| PC5 | 3.7 | 11 | 3.217 | 3.015 | 0.002 |
| PC6 | 3.1 | 11 | 3.249 | 3.298 | 0.001 |
| PC7 | 2.7 | 11 | 1.653 | 1.292 | 0.093 |
| PC8 | 2.4 | 11 | 1.832 | 1.598 | 0.046 |
| PC9 | 2.0 | 11 | 3.244 | 3.175 | 0.003 |
| ***plod2* crispant vs control** | | | | | |
| PC1 | 19.8 | 1 | 0.041 | -1.029 | 0.831 |
| PC2 | 13.5 | 1 | 7.886 | 2.403 | 0.003 |
| PC3 | 11.6 | 1 | 4.224 | 1.519 | 0.066 |
| PC4 | 7.3 | 1 | 1.425 | 0.728 | 0.245 |
| PC5 | 6.3 | 1 | 0.780 | 0.311 | 0.395 |
| PC6 | 5.8 | 1 | 0.021 | -1.304 | 0.892 |
| PC7 | 5.0 | 1 | 0.005 | -1.664 | 0.938 |
| PC8 | 4.4 | 1 | 0.850 | 0.414 | 0.346 |
| PC9 | 3.7 | 1 | 0.005 | -1.554 | 0.925 |
| PC10 | 3.6 | 1 | 2.815 | 1.293 | 0.105 |
| PC11 | 2.9 | 1 | 0.403 | -0.047 | 0.524 |
| PC12 | 2.8 | 1 | 1.737 | 0.904 | 0.181 |
| PC13 | 2.5 | 1 | 0.024 | -1.230 | 0.865 |
| PC14 | 2.5 | 1 | 0.096 | -0.776 | 0.760 |
| PC15 | 2.1 | 1 | 0.074 | -0.819 | 0.776 |
| ***meox1* crispant vs control** | | | | | |
| PC1 | 28.9 | 1 | 4.977 | 1.686 | 0.048 |
| PC2 | 13.2 | 1 | 2.271 | 1.085 | 0.131 |
| PC3 | 11.1 | 1 | 8.981 | 2.292 | 0.012 |
| PC4 | 8.0 | 1 | 0.003 | -1.715 | 0.953 |
| PC5 | 6.0 | 1 | 0.069 | -0.936 | 0.797 |
| PC6 | 5.2 | 1 | 1.628 | 0.767 | 0.239 |
| PC7 | 4.1 | 1 | 0.350 | -0.096 | 0.538 |
| PC8 | 3.5 | 1 | 0.713 | 0.305 | 0.394 |
| PC9 | 3.2 | 1 | 0.042 | -1.001 | 0.828 |
| PC10 | 2.5 | 1 | 0.007 | -1.579 | 0.932 |
| PC11 | 2.1 | 1 | 0.093 | -0.757 | 0.760 |
| ***sost* mutant vs control** | | | | | |

| | | | | | |
|---|---|---|---|---|---|
| PC1 | 38.0 | 1 | 0.013 | -1.411 | 0.916 |
| PC2 | 20.4 | 1 | 17.045 | 2.787 | 0.002 |
| PC3 | 8.9 | 1 | 0.176 | -0.478 | 0.683 |
| PC4 | 6.1 | 1 | 0.025 | -1.233 | 0.882 |
| PC5 | 5.4 | 1 | 1.097 | 0.563 | 0.305 |
| PC6 | 4.3 | 1 | 0.102 | -0.711 | 0.761 |
| PC7 | 3.7 | 1 | 0.336 | -0.139 | 0.561 |
| PC8 | 3.2 | 1 | 0.136 | -0.603 | 0.728 |
| PC9 | 2.9 | 1 | 0.040 | -1.250 | 0.874 |

**Table S4.** Procrustes ANOVA results for analyses of symmetric components of shape variation in this study. The groupings test if PCs differ between crispant/mutant and controls (PC ~ group). The proportion of variance explained by each PC is given in the %Var column.

| PC | % Var | DF | F | Z | p |
|---|---|---|---|---|---|
| ***plod2* crispant vs control** | | | | | |
| PC1 | 27.3 | 1 | 0.140 | -0.615 | 0.722 |
| PC2 | 15.7 | 1 | 16.587 | 2.813 | 0.001 |
| PC3 | 9.7 | 1 | 1.261 | 0.644 | 0.263 |
| PC4 | 8.5 | 1 | 1.484 | 0.808 | 0.234 |
| PC5 | 5.9 | 1 | 0.009 | -1.546 | 0.936 |
| PC6 | 5.3 | 1 | 0.304 | -0.168 | 0.579 |
| PC7 | 3.9 | 1 | 0.374 | -0.125 | 0.547 |
| PC8 | 3.5 | 1 | 3.276 | 1.355 | 0.081 |
| PC9 | 3.2 | 1 | 0.297 | -0.216 | 0.595 |
| PC10 | 2.7 | 1 | 0.005 | -1.578 | 0.935 |
| PC11 | 2.5 | 1 | 0.532 | 0.151 | 0.462 |
| PC12 | 2.1 | 1 | 0.670 | 0.179 | 0.459 |
| ***meox1* crispant vs control** | | | | | |
| PC1 | 41.8 | 1 | 2.632 | 1.181 | 0.123 |
| PC2 | 17.0 | 1 | 0.007 | -1.584 | 0.939 |
| PC3 | 10.0 | 1 | 5.636 | 1.849 | 0.029 |
| PC4 | 5.3 | 1 | 1.694 | 0.886 | 0.197 |
| PC5 | 4.2 | 1 | 2.421 | 1.097 | 0.140 |
| PC6 | 3.6 | 1 | 0.626 | 0.214 | 0.443 |
| PC7 | 3.3 | 1 | 1.708 | 0.837 | 0.210 |
| PC8 | 2.6 | 1 | 0.023 | -1.292 | 0.883 |
| PC9 | 2.0 | 1 | 0.626 | 0.252 | 0.420 |
| ***sost* mutant vs heterozygote vs control** | | | | | |
| PC1 | 34.2 | 2 | 8.147 | 2.709 | 0.005 |
| PC2 | 13.3 | 2 | 0.106 | -1.316 | 0.897 |
| PC3 | 11.4 | 2 | 0.075 | -1.437 | 0.914 |
| PC4 | 9.2 | 2 | 0.368 | -0.449 | 0.683 |
| PC5 | 5.8 | 2 | 0.188 | -0.994 | 0.830 |
| PC6 | 4.0 | 2 | 0.986 | 0.278 | 0.398 |
| PC7 | 3.4 | 2 | 0.957 | 0.266 | 0.406 |
| PC8 | 3.0 | 2 | 0.354 | -0.517 | 0.698 |
| PC9 | 2.3 | 2 | 0.214 | -0.910 | 0.817 |
| PC10 | 2.2 | 2 | 2.864 | 1.394 | 0.075 |
| ***wnt16* crispant vs control** | | | | | |
| PC1 | 41.6 | 1 | 3.355 | 1.415 | 0.073 |
| PC2 | 11.5 | 1 | 10.173 | 2.542 | 0.002 |
| PC3 | 8.5 | 1 | 2.637 | 1.215 | 0.107 |
| PC4 | 5.9 | 1 | 0.080 | -0.824 | 0.790 |

| | | | | | |
|---|---|---|---|---|---|
| PC5 | 5.4 | 1 | 1.573 | 0.820 | 0.210 |
| PC6 | 3.9 | 1 | 0.264 | -0.273 | 0.621 |
| PC7 | 3.1 | 1 | 0.033 | -1.233 | 0.874 |
| PC8 | 2.8 | 1 | 0.035 | -1.059 | 0.839 |
| PC9 | 2.5 | 1 | 0.703 | 0.274 | 0.410 |

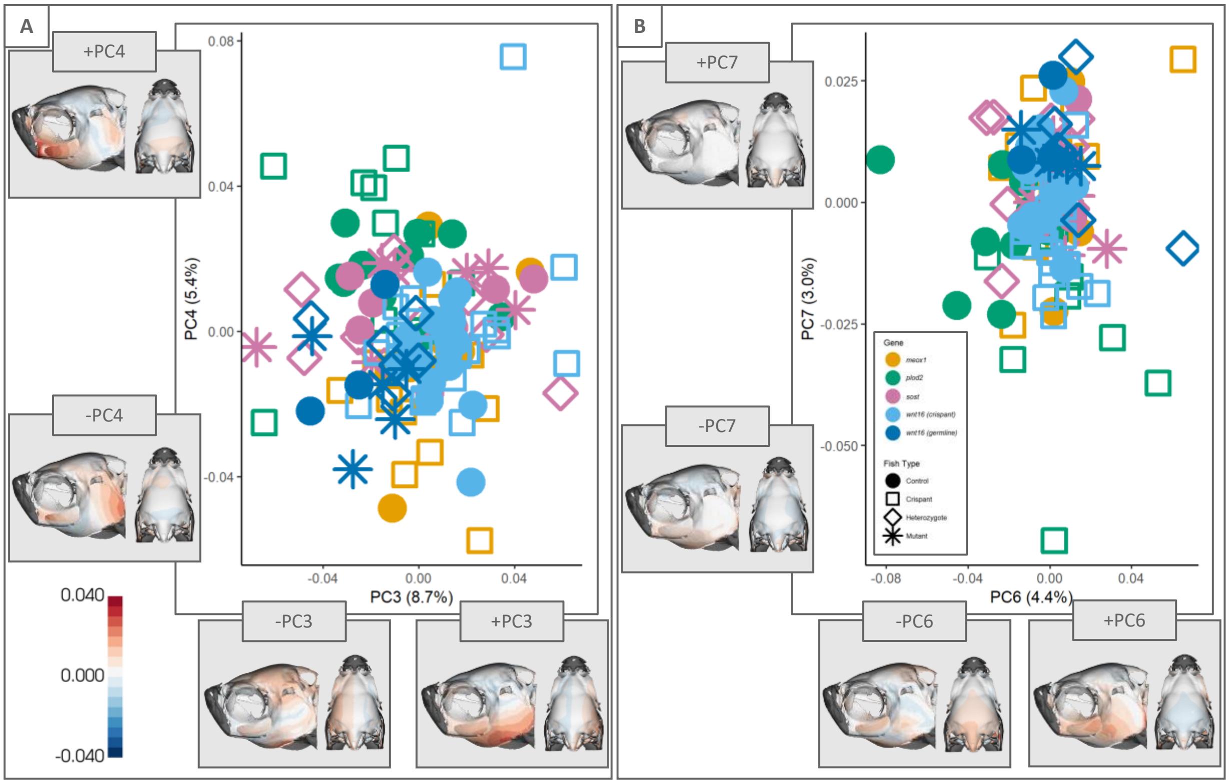


**Figure S1.** Additional principal components showing groupwise difference from analyses using manual landmarks. Point color indicates gene grouping; shape indicates type of fish. Heatmaps represent lateral (left) and dorsal (right) cranial skeletons and show Procrustes distances between mean shape and the extreme shapes across each PC. All axis labels show the percentage of variation represented by each PC.

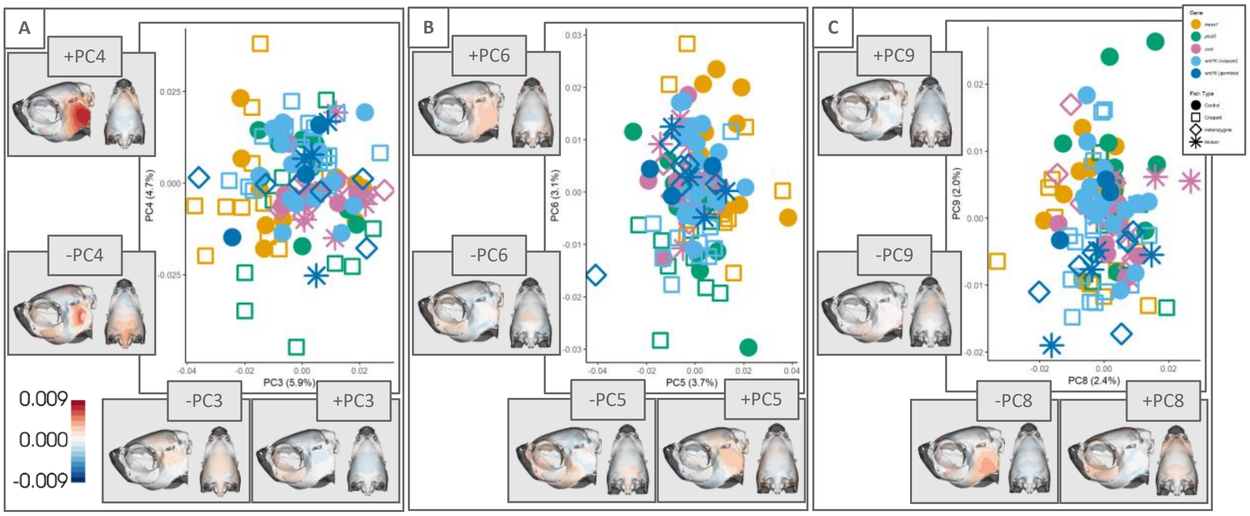


**Figure S2.** Additional principal components showing groupwise difference from analyses using pseudolandmarks. Point color indicates gene grouping; shape indicates type of fish. Heatmaps represent lateral (left) and dorsal (right) cranial skeletons and show Procrustes distances between mean shape and the extreme shapes across each PC. All axis labels show the percentage of variation represented by each PC.

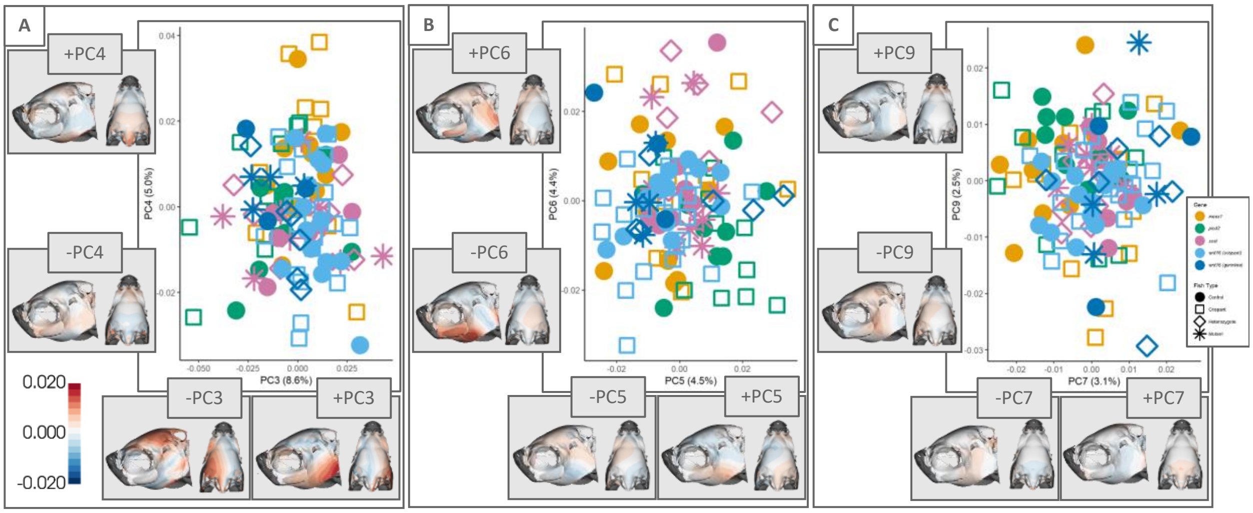


**Figure S3.** Additional principal components showing groupwise difference from analyses using the MALPACA placed landmarks. Point color indicates gene grouping; shape indicates type of fish. Heatmaps represent lateral (left) and dorsal (right) cranial skeletons and show Procrustes distances between mean shape and the extreme shapes across each PC. All axis labels show the percentage of variation represented by each PC.

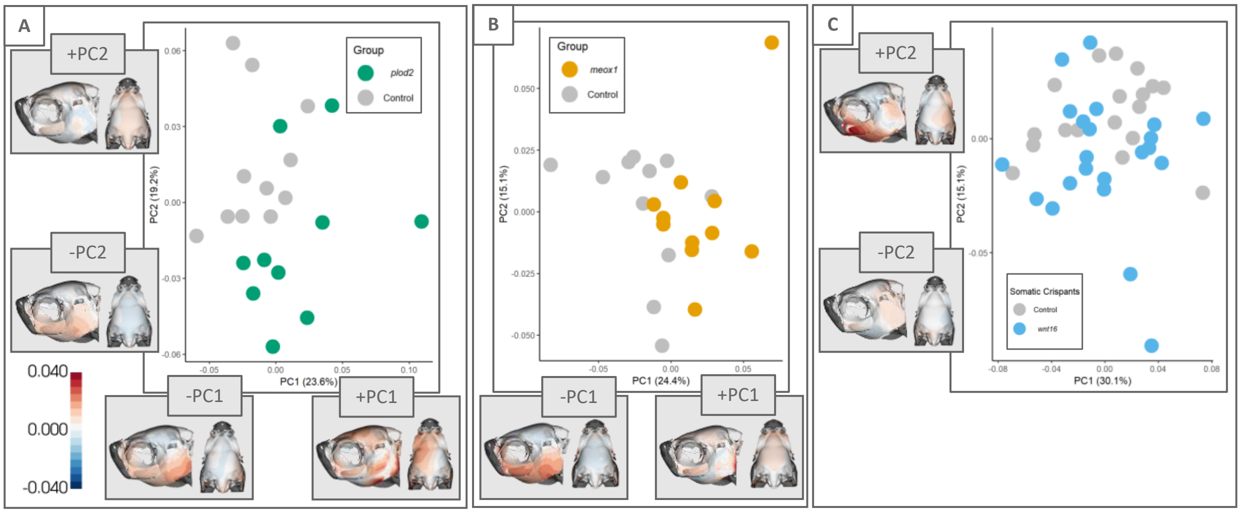


**Figure S4.** Principal component plots from manual landmark analyses that show separation of crispant and control fish for A) *plod2*, (B) *mox1*, and (C) *wnt16* crispants. Point color indicates groupings. Heatmaps represent lateral (left) and dorsal (right) cranial skeletons and show Procrustes distances between mean shape and the extreme shapes across each PC. Axis labels show the percentage of variation represented by each PC.

**Appendix 1:** R script showing an example of our statistical analysis for a single gene. (Link to script on GitHub will be added after acceptance of manuscript, file added as supplemental information for submission).

**Appendix 2:** ZFMask.py Python script for creating heatmaps of mesh distances in 3D Slicer. (Link to script on GitHub will be added after acceptance of manuscript, file added as supplemental information for submission).
